# Distinct neural routes for evaluative and informational social influence

**DOI:** 10.1101/2025.10.30.685641

**Authors:** Sriranjani Manivasagam, Thomas Schultze, Anne Schacht, Anna Fischer

**Affiliations:** University of Göttingen, Institute of Psychology, Germany; University of Bamberg, Institute of Psychology, Germany; Queen’s University Belfast, School of Psychology, United Kingdom

**Author notes:** **Corresponding author:** (AS). **Author note:** Anna Fischer and Anne Schacht contributed equally to this work and share senior authorship.

**Keywords:** Social influence, Advice taking, Feedback, Metacognition, Confidence, Electroencephalography, Judgment and decision-making

## Abstract

Humans routinely revise their judgments based on input from others, yet how the brain mechanisms distinguish evaluative from informational social influence remains unclear. Using preregistered electroencephalography (EEG, N = 87) and the Judge-Advisor System, we identified distinct neurocognitive routes by which feedback and advice shape judgment revision. Participants made numerical estimates, received either feedback (“good”/“bad”) or advice (alternative estimates) of varying proximity to their own judgment, and then revised their response.

Feedback engaged rapid evaluative monitoring: a feedback-related negativity (FRN, 200-300ms) that decreased with confidence, indicating reflexive performance assessment. Advice triggered delayed deliberate processing: late positive complex (LPC) onset latencies that slowed when confident individuals encountered discrepant advice, indicating effortful integration of external information. Both routes converged on the P300 (250-250ms),whose amplitudes increased with proximity, consistent with the reward value of social validation.

These findings reveal a dual-route architecture: feedback determines *whether* to revise through rapid evaluative signals, while advice specifies *how much* to revise through sustained integrative processes. Confidence calibrates both routes, protecting against misleading feedback while intensifying deliberation over discrepant advice. This temporal dissociation clarifies how metacognitive and social processes coordinate adaptive judgment revision.

## Introduction

Human decisions are rarely made in isolation. Social information such as advice, feedback, or consensus cues can substantially alter confidence and choice behavior. While the motivational and informational value of social cues has been widely studied, the neural mechanisms that determine how individuals evaluate and integrate such information remain unclear. Understanding these mechanisms is essential, as social information can both enhance judgment accuracy and introduce systematic biases in decision making.

The distinction between evaluative and informational influence has broad practical implications. In educational settings, feedback shapes learning trajectories but can undermine motivation when delivered poorly (Hattie & Timperley, 2007; Kluger & DeNisi, 1996), while concrete guidance enables targeted skill improvement. In organizational contexts, advice-taking determines whether groups leverage diverse perspectives or succumb to premature consensus (Kämmer et al., 2023; Yaniv, 2004). In human-AI collaboration, systems must balance providing evaluative confidence scores with actionable recommendations to optimally support users. Yet despite decades of behavioral research documenting when people accept or reject social influence, the neurocognitive processes that determine how individuals distinguish evaluative from informational input – and integrate each type into their own judgments – remain largely uncharacterized. Identifying these mechanisms could inform evidence-based practices for optimizing social influence across domains, from education to collective decision-making to human-machine interaction.

Integrating social and personal information allows individuals to optimize decision accuracy, and the extent of this integration depends on confidence calibration and metacognitive control (Bahrami et al., 2010). Confidence not only predicts whether individuals will adjust their behavior in response to social input but also modulates the neural mechanisms of monitoring and control (Boldt & Yeung, 2015; Boldt et al., 2017). High confidence typically reduces susceptibility to external influence, whereas low confidence increases the likelihood of adopting others’ opinions. Yet, confidence interacts with contextual factors – such as the perceived proximity or similarity of social information – to determine when and how individuals integrate external advice or feedback into their own judgments (Pescetelli & Yeung, 2021).

Research on social influence distinguishes two motivational routes by which others shape behavior (Deutsch & Gerard, 1955). Normative influence arises from the affiliation motive and leads individuals to align their publicly stated opinions with others to avoid social exclusion. Informational influence, in contrast, reflects an epistemic motive: individuals revise their judgments because they accept information from others as evidence about reality. The present study focuses on this latter form of influence, which has been systematically examined using the Judge-Advisor System (JAS) (Sniezek and Buckley,1995), in which individuals (the *judges*) make an initial estimate, receive social information from another person (the *advisor*), and then decide whether to revise it. In the classic JAS, judge and advisor do not interact, minimizing normative social influence. Prior work has predominantly focused on behavioral outcomes (Kämmer et al., 2023), leaving the underlying neurocognitive mechanisms of informational social influence largely unexplored.

Advice can take different forms (Dalal & Bonaccio, 2010), two of which we are interested here. One type involves evaluative recommendations that support or discourage a particular course of action (Dalal & Bonaccio, 2006). We define these statements – such as indicating that an estimate is “good” or “bad” – as Feedback, a binary evaluation that reinforces or challenges an individual’s initial judgment. Another type, informational advice, goes beyond simple evaluation to provide concrete suggestions for adjustment, such as alternative numerical estimates, usually in the form of another person’s explicit judgment or decision (Bonaccio & Dalal, 2006; Schultze et al., 2015). In line with the advice-taking literature, we simply refer to this type of specific recommendation as Advice.

Feedback and advice both serve to reduce uncertainty but differ in their informational content and cognitive demands. Feedback provides an explicit evaluation (“good” or “bad”) that prompts rapid self-assessment but offers no direction for change. Advice, in contrast, conveys specific alternative information, but this information requires interpretation and, potentially, integration with one’s prior judgment. Consider for example a person who discusses with a colleague how much to offer in an upcoming negotiation. This person might ponder offering an amount of $50,000. The colleague might comment that that is a “bad idea” or that it “sounds reasonable” (feedback). However, the colleague could also propose a specific alternative by stating “I would offer $52,000” (advice). In the latter case, the decision-maker must first interpret the advice to

determine whether the original offer of $50,000 is good enough in the light of the numerically different advice. In case the decision-maker decides that it is not, the next question is how to adjust the offer, for example by increasing it to $51,000. Put simply, where feedback is unequivocal in its evaluative statement, advice still contains an evaluative component that is open to interpretation. However, advice also contains specific informational value while feedback is ambiguous in that regard. Importantly, these two components of social information may engage separable neural routes: an *evaluative route* that monitors performance and an *informational route* that supports deliberative adjustment. The present study tests this distinction directly.

Informational social influence depends on how close or discrepant the social information is relative to one’s own initial judgment (Yaniv, 2004a; Yaniv & Milyavsky, 2007; Minson et al., 2011; Moussaïd et al., 2013; Schultze et al., 2015; Du et al., 2019) – a property we refer to as proximity. Proximity captures the extent to which external input aligns with an individual’s estimate and thereby determines its motivational and informational value, which in turn interacts with confidence to shape social influence. In this framework, the proximity of advice mirrors the valence of feedback: high-proximity advice, like positive feedback, signals that one’s judgment is close to a desired or correct outcome and thus reinforces confidence, whereas low-proximity advice, like negative feedback, indicates a larger discrepancy and challenges the initial judgment. For consistency, we use the term *proximity* throughout to denote the gap between individuals’ initial estimates and the social information provided, irrespective of whether that information is feedback or advice.

Event-related potentials (ERPs) provide the temporal precision needed to distinguish early evaluative from later integrative stages of social information processing. The feedback-related negativity (FRN), emerging around 200–300 ms post-stimulus at frontocentral sites, indexes rapid outcome evaluation and performance monitoring (Miltner et al., 1997; Yeung et al., 2004; Yeung & Sanfey, 2004; Holroyd et al., 2006; Goyer et al., 2008; San Martín, 2012). The P300 (250–350 ms, centroparietal) reflects attention allocation and the updating of motivationally significant information (Sutton et al., 1965; Donchin & Coles, 1998; Polich, 2007; Nieuwenhuis et al., 2005). Similar to the FRN, the P300 has also been demonstrated to be sensitive to both the magnitude and valence of outcomes (Yeung & Sanfey, 2004; Hajcak et al., 2005; Sato et al., 2005; Wu & Zhou, 2009). The late positive complex (LPC) (≈400–600 ms and beyond) marks sustained elaboration and integration of task-relevant content (Polich, 2007; Fields, 2023) and is sensitive to stimulus salience and motivational relevance (Cuthbert et al., 2000; Schacht & Sommer, 2009; Bayer & Schacht, 2014; Hajcak et al., 2010; Bayer & Schacht, 2014; Grassi et al., 2023; Ziereis & Schacht, 2024). Recent work using social learning paradigms further refines this view, demonstrating that the FRN predominantly indexes *reward* prediction errors, whereas later parietal positivities (P3b/LPP) track *affective* prediction errors (Heffner et al., 2025). This neural separability between reward and emotion learning signals provides converging evidence for temporally distinct evaluative and integrative processes in social information use. Together, these components trace the transition from reflexive evaluation to deliberate revision, making them ideally suited to dissociate neural routes engaged by feedback and advice. However, these neural responses are not uniform across individuals or contexts. Their magnitude and timing are shaped by metacognitive factors, particularly by how confident individuals are in their initial judgments – a variable known to determine how strongly people rely on social input.

Confidence modulates informational social influence by determining how extensively people rely on external input for subsequent judgments (Desender et al., 2018; Pescetelli & Yeung, 2021; Frömer et al., 2021; Carlebach & Yeung, 2023). Serving as a metacognitive indicator of certainty, confidence guides information-seeking: when individuals are uncertain, they place greater weight on others’ input; when confident, they rely more on their own judgment. Importantly, confidence interacts with proximity. Low proximity is attributed to decision-makers’ own initial opinion being wrong when confidence is low, but to the advice or feedback being wrong when confidence is high (Pescetelli & Yeung, 2021). This calibration enables individuals to assess the reliability of social information and its source, improving decision outcomes.

Building on this framework, the present preregistered study examined how advice and feedback – depending on their proximity to individuals’ initial estimates – shape judgment and decision-making. This comparison allowed us to identify the neurocognitive mechanisms shared across both types of social information and those specific to advice’s additional informational value. Because we did not predict directional differences between feedback and advice, our hypotheses focused on the effects of social information proximity and its interaction with confidence. Behaviorally, we expected greater influence from low-proximity (discrepant) than from high-proximity (confirming) social information, and higher post-information confidence following high-proximity input. For the ERP components of interest, we predicted that low-proximity information would elicit enhanced feedback-related negativity (FRN) amplitudes, reflecting stronger performance monitoring, and increased late positive complex (LPC) amplitudes, indicating greater cognitive resources for updating judgments. Conversely, high-proximity information was expected to evoke larger P300 amplitudes, consistent with the reward value of social validation. Finally, we exploratively examined how individuals’ initial confidence modulated these neural and behavioral effects.

## Methods

The study was preregistered on the Open Science Framework (OSF) before data collection (https://osf.io/nvfh5).

### Participants

We recruited 110 participants (mean age = 22.42 ± 3.34 years; 89 female, 20 male, 1 other; 108 right-handed, 2 left-handed) through a combination of online and offline advertisements, flyers, mailing lists, and social media. All participants were native German speakers with normal or corrected-to-normal vision and no neurological or psychiatric history. Participants received 8.50 EUR per hour or course credits, plus a 25 EUR bonus for performance within the top 20 % of all participants, thus providing an additional incentive for optimal task performance. All participants had the opportunity to clarify questions after the experimental procedure was explained to them and gave written informed consent before participating in the experiment. The study was approved by the ethics committee of the Georg-Elias-Mueller-Institute for Psychology of the University of Goettingen (application number: 319, 2022-11-11) and conducted in accordance with the Declaration of Helsinki.

As preregistered, participants with excessive EEG artifacts were excluded and replaced. Due to unexpectedly high artifact rates during summer recordings, the exclusion threshold for trial loss was increased from 20% to 30% before data analysis and applied uniformly. Data from 20 participants were excluded because of excessive artifacts; one additional dataset was removed due to corrupted trigger codes, and two left-handed participants were excluded to maintain a right-handed sample. The final sample comprised 87 participants (mean age = 22.38 ± 3.49 years; 72 female, 15 male; all right-handed).

### Stimuli and Design

Stimuli consisted of estimation questions about (1) the aerial distance between two European cities and (2) the caloric content of various food items (adapted from Schultze et al., 2015). The experiment followed a 2 x 2 within-subject design with the factors social information type (advice, feedback) and social information proximity (high, low). Each participant completed two task blocks—one for distance and one for calorie estimations—while receiving either advice or feedback. The assignment of information type to task block was counterbalanced across participants. Each block contained 100 randomized trials (50 high- and 50 low-proximity).

In the *advice* condition, participants received numerical suggestions derived from their own initial estimates. High-proximity advice was created by adding or subtracting a random value between 8–12%, and low-proximity advice by adding or subtracting 48–52% from the participant’s estimate. Values were rounded to tens to enhance plausibility. In the *feedback* condition, participants received the verbal statements “good” (high proximity) or “bad” (low proximity), each appearing with equal probability and randomly assigned to trials irrespective of accuracy. This ensured that feedback valence was orthogonal to objective performance and directly comparable to advice proximity.

To prevent implausible social information when participants entered obviously incorrect initial estimates (e.g., 15 km instead of 150 km), a failsafe mechanism was implemented. When initial estimates fell outside predefined ranges (25–7.500 km for distance; 10–2.000 kcal for calories), advice was generated relative to the true value, and feedback defaulted to *“bad.”* Trials in which the failsafe mechanism was triggered were excluded from all behavioral and EEG analyses. In addition, trials without a final estimate or confidence rating were excluded from behavioral analyses.

### Procedure

Each trial followed the sequence illustrated in Figure 1. An estimation question appeared and remained on screen until response (maximum 15 s). Participants entered an initial estimate and rated their confidence on a 7-point Likert scale (1 = not at all confident; 7 = very confident; maximum 10 s). Next, participants received social information – either advice or feedback – of high or low proximity for 1 s. In advice trials, this consisted of the computed alternative estimate; in feedback trials, of the verbal label *“good”* or *“bad”.* After receiving social information, participants had the opportunity to revise their estimate and then provided a final confidence rating on the same scale. All trial types (advice/feedback × high/low proximity) occurred with equal frequency and in fully randomized order within each block.

**Figure 1.**
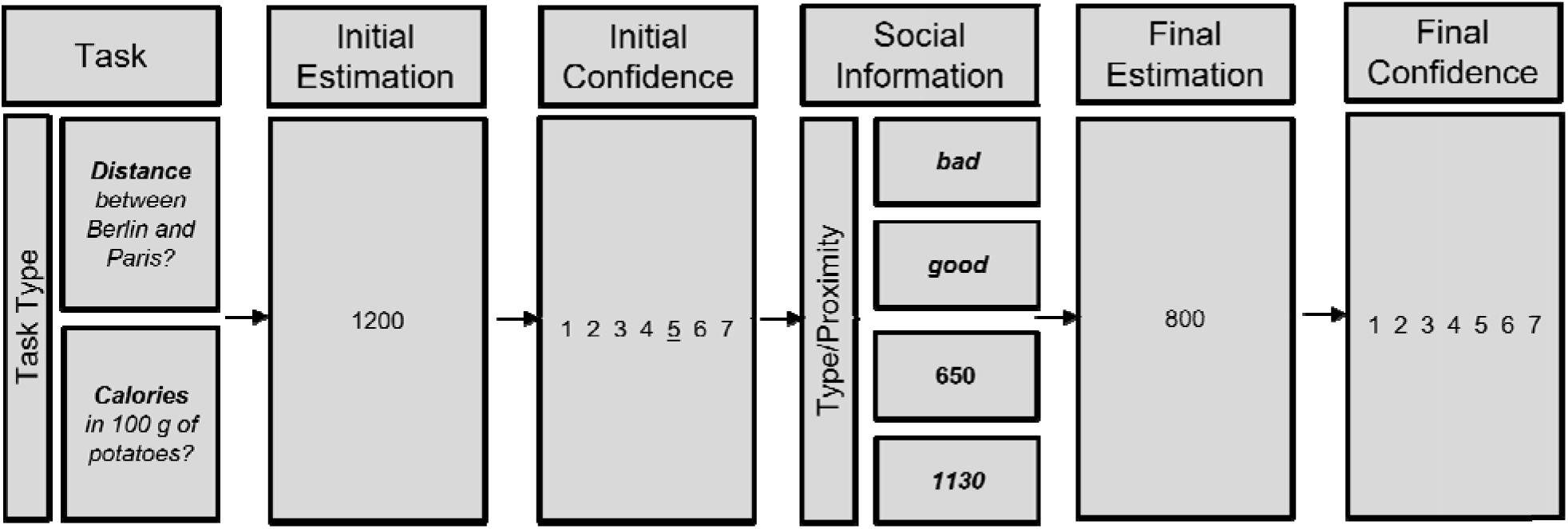
*Estimation Tasks using the Judge-Advisor-System (JAS).* Participants performed two blocks, one of each task type: distance and calories. Within each block, they consistently received one of two types of social information – ‘advice’ or ‘feedback’. The assignment of social information type to blocks was counterbalanced across participants. In each trial, participants were presented with an estimation question, which remained on the screen until they provided their response (maximum 15 seconds). They then made their initial estimation, followed by a confidence rating on a 7-point scale (1 – not at all confident to 7 – very confident; maximum 10 seconds for response). Next, they received social information (‘advice’ or ‘feedback’) of either high or low proximity, presented for 1 second with equal probability. Participants then had the opportunity to revise their decision by making a final estimation, followed by a final confidence rating.

### Behavioral Measures

Within the JAS (see Figure 1), we recorded participants’ initial (before social information) and final (after social information) estimates, as well as their confidence ratings for both estimates on a 7-point Likert scale (1 = not at all confident; 7 = very confident. Response times for all estimates and confidence ratings were logged at the first button press. From the behavioral data, the following measures were extracted and calculated as follows:

*Decision to Adjust*: This binary variable captured whether participants revised their estimates after receiving social information. Trials in which participants either refrained from providing a revised estimate or repeated their initial value were coded as 0; all other trials were coded as 1.

*Relative Decision Change (AT Score)*: AT Score (Harvey & Fischer, 1997), which takes the percent weightage given to advice into account. The ‘AT Score’ is defined as follows:

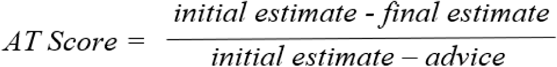

The AT Score was computed only for advice trials and winsorized to the range [0,1] following field standards (Soll & Larrick, 2009; Schultze et al., 2015), such that values > 1 were set to 1, and values < 0 were set 0. A total of 10.43 % of trials were winsorized.

*Confidence Shift*: This variable reflected the difference between the participants’ confidence final and initial confidence ratings. Positive values indicated increased confidence after receiving social information, whereas negative values indicated decreased confidence.

### EEG Recording and Preprocessing

The EEG was recorded during the experiment from 128 active Ag/AgCl electrodes positioned in an electrode cap (Easy Cap) that allows attachment of the electrodes according to the extended 10–20 system (Pivik et al., 1993). External electrodes were attached bilaterally on the mastoids, the outer canthi (HEOG) and below the eyes (VEOG). The common mode sense (CMS) active electrode was used as reference, and the driven right leg (DRL) passive electrode served as ground. A Biosemi ActiveTwo AD-Box (24 bits; band-pass filter 0.16–100 Hz) was used to amplify the signal recorded by the ActiView software at a sampling rate of 512 Hz. Electrodes offsets were kept between +/–20 mV.

The raw EEG data were preprocessed using the EEGLAB (version 2022.0; Delorme & Makeig, 2004), ERPLAB (version 9.10; Lopez-Calderon & Luck, 2014), and BioSig (version 3.8.1; Vidaurre et al., 2011) toolboxes in MATLAB (R2018a). The raw signal was referenced to the average of mastoids and filtered offline with a 0.1 Hz high-pass and a 40 Hz low-pass filter. The data were segmented from −200 to 1000 ms relative to the onset of the social information presentation and baseline-corrected to a 200 ms pre-stimulus interval. Independent Component Analysis (ICA) was run using the *runica* algorithm of the EEGLAB toolbox to remove artifacts such as eye blinks, muscle activity and channel noise by manual inspection of the components. The data were visually inspected to identify channels with poor signal quality; any removed channels were interpolated using the spherical interpolation algorithm. Subsequently, trials exhibiting excessive noise were excluded if voltage values exceed the thresholds of ± 100 µV within the time window of −200 ms to 700 ms, or if the slope of the voltage trend reached 100 µV or more within the ERP analysis window. Further, data were defined as statistically improbable and excluded if values in a trial exceeded 5 SD of the mean probability distribution, both at the individual channel level as well as for all channels combined.

Finally, the signal was re-referenced to the average of all channels for further statistical analysis. ERP measures extracted were mean FRN, P300, and LPC amplitudes for each trial, time-locked to the onset of social information: a) FRN, frontocentral electrode cluster (channels c11, c12, c21, c22, c23, c24, and c25), 200-300 ms; P300, centroparietal electrode cluster (channels a02, a03, a04, a05, a19, and a32), 250-350 ms; LPC, same electrode cluster as P300, 400-600 ms.

The onset latencies of these ERP components were calculated using a relative criterion technique, identifying the time point at which the ERP amplitude reached 50 % of its maximum (P300, LPC) or minimum (FRN) within the respective time windows (Kiesel et al., 2008). Because we employed mixed models using single trial onset latencies per participant, we did not employ the jackknife-based approach recommended by the same authors in conjunction with this technique.

### Overview of the statistical workflow

All statistical analyses and visualization were conducted in RStudio (RStudio version 2024.04.2; R version 4.3.2). To statistically test all the predicted effects, linear mixed models (LMM) with a gaussian error distribution and ‘identity’ link function or generalized linear mixed models (GLMM) with a binomial error structure and ‘logit’ link function with maximal random effects structures were employed by including all theoretically identifiable random slopes and their interactions (Schielzeth & Forstmeier, 2009; Barr et al., 2013) using the function ‘lmer’ and ‘glmer’ of the package lme4 (version 1.1-29; Bates et al., 2015), respectively. For all the models of interest, subject ID was used as the grouping factor in the random effects part and correlations between random intercepts and slopes were included. The ‘Decision to Adjust’ variable was modeled using a GLMM, given its binomial response structure, whereas all other models were LMMs. For the factors ‘social information type’ and ‘social information proximity’, the levels ‘advice’ and ‘high’, respectively, were used as the reference levels in all models. An exception was made for the models predicting P300 amplitudes, where the level ‘low’ was used as the reference for ‘social information type’ in order to align the model estimates with the directional hypotheses.

The analyses we report here deviate from the preregistered analyses in two ways: First, we included judges’ confidence in the initial estimates (z-transformed within participants) as a covariate in all models except the behavioral model for ‘Confidence Shift’. This addition was theoretically motivated by Pescetelli and Yeung’s (2021) work which we became aware of only after we ran our study. As stated above, this work indicates that proximity is not necessarily informative on its own in social information processing and instead must be considered contingent on decision-makers’ confidence. Second, we included trial number as a covariate in the fixed-effects structure of all the models due to methodological considerations. Since our study contained a large number of similar estimation tasks, including trial number allows us to capture possible effects such as loss of motivation over time on the behavioral level or task-related habitual effects for the ERPs. The covariate trial number was z-transformed to improve model convergence and enhance the interpretability of the model intercept. Incorporating trial number allowed us to retain the full granularity of trial-level variability and provided more data points for estimating model parameters, thereby increasing statistical power (Aarts et al., 2014). However, to avoid overestimating the precision of fixed-effects estimates and to maintain the Type I error rate at the nominal 5 % level, the trial number was also included as a random slope within subject

ID (Schielzeth & Forstmeier, 2009; Barr et al., 2013). Furthermore, to assess the contribution of the fixed effects of interest beyond chance, we performed full-null model comparisons (Forstmeier & Schielzeth, 2011), where the null models excluded the predictors of interest but retained the control variable (trial number) and identical random-effects structures as the full models. For full transparency, we report a full overview of the preregistered analyses in the supplement. In brief, the addition of the two predictors yielded two main insights: first, there were trial effects on the behavioral level, but not on the ERP data, and including trial-level effects on the behavioral level did not change any of the results qualitatively. Second, adding confidence did not change any results on the behavioral level, but revealed strong interaction effects in some of the ERP analyses that would have otherwise gone unnoticed.

The contribution of each predictor was assessed using a likelihood ratio test (Dobson & Barnett, 2018) comparing all possible models by dropping single terms using the function ‘drop1’. The function systematically removes one predictor at a time from the full model and evaluates whether its exclusion significantly worsens model fit, thereby indicating its importance. For the LMMs, restricted maximum likelihood was used for refitting the models to obtain p-values by means of the Satterthwaite approximation (Luke, 2017) with the function lmer of the package lmerTest (version 3.1-3; (Luke, 2017). For the GLMM model, p-values were obtained from the ‘glmer’ of the package lme4 (version 1.1-29; Bates et al., 2015).

For the LMM models, we checked the assumptions of normality and homogeneity of the residuals by means of qq-plots and residuals plotted against fitted values. We further assessed the model stability using the DFBeta method by successively excluding levels of the grouping factor to ascertain the effects on the model coefficients.

In the instance that the full-null model comparison revealed significance, while the three-way interaction between social information type, social information proximity, and initial confidence did not, we eliminated the three-way interaction, and subsequently any potential non-significant two-way interactions, to obtain reliable estimates that are unconditional on particular values of other predictors with which they interact in the model. Finally, we obtained 95 % confidence intervals for the model estimates through parametric bootstrapping (N = 1000). For interactions, post hoc tests were conducted using the ’emmeans’ package (version 1.10.2; (Lenth, 2020)) and p-values were adjusted using ‘Tukeys’ method to keep the chance of Type-1 error to a minimum.

## Results

### Behavioral data

Across all participants, 539 trials (3.14%) activated the failsafe mechanism. The final dataset for behavioral analysis comprised 16,560 (advice–high proximity = 4,028; advice–low proximity = 4,099; feedback–high proximity = 4,205; feedback–low proximity = 4,228).

***Decision to Adjust*** *(Figure 2A).* The binary *Decision to Adjust* variable (0 = no change, 1 = revised estimate) was significantly modulated by an interaction between social information proximity and information type (proximity x type: β = 2.697, SE = 0.270, z = 9.975, p < .001). Participants were more likely to adjust their estimates after receiving low-compared with high-proximity advice (β = 2.21, SE = 0.127, z = 17.340, p < .001). This proximity effect was even stronger for feedback (β = 4.91, SE = 0.227, z = 21.616, p < .001). Additionally, adjustment probability decreased with increasing initial confidence (β = –0.277, SE = 0.036, t = –7.718, p < .001; *Figure 3A*).

**Figure 2.**
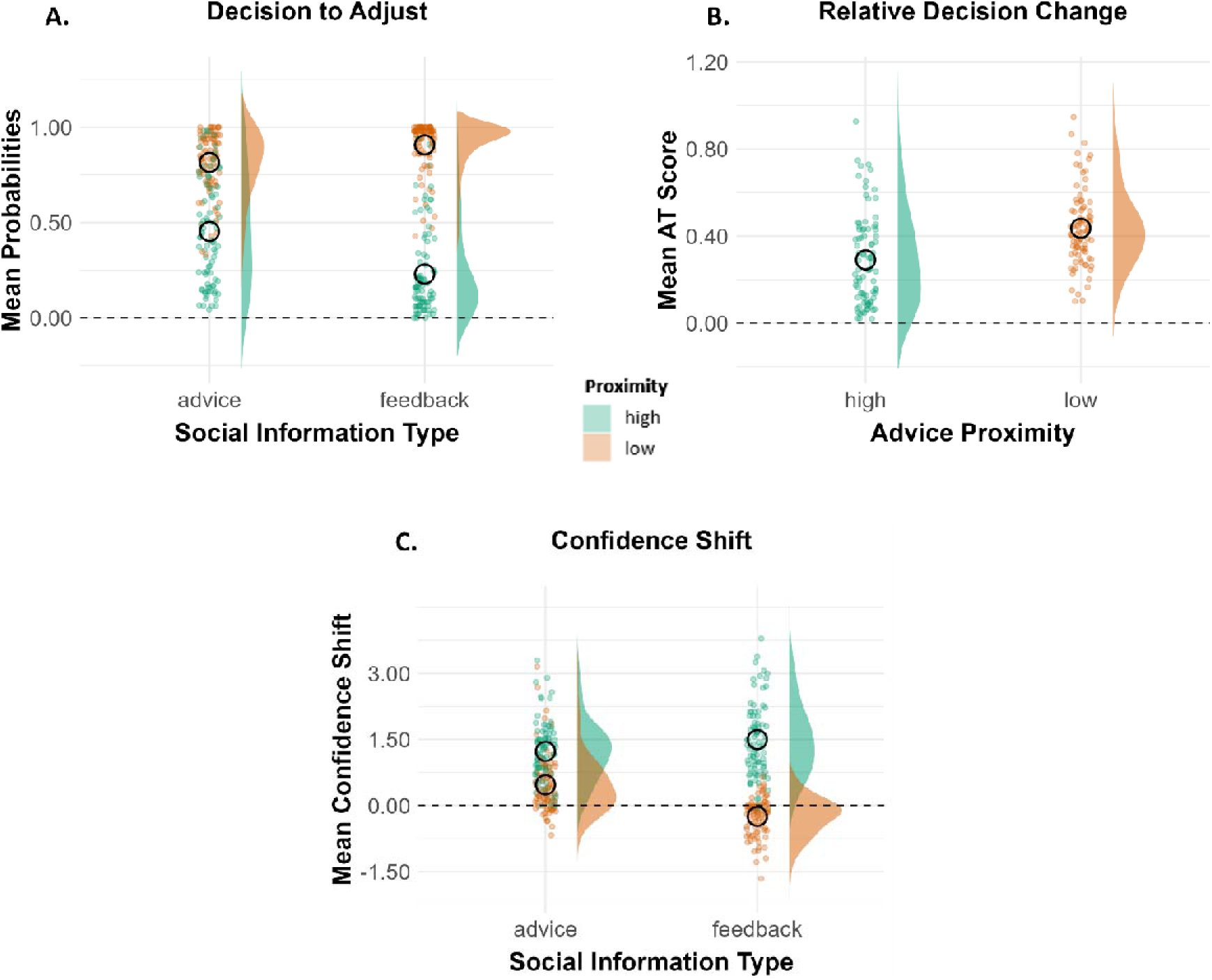
***A.*** *Decision to Adjust **B.** Relative Decision Change (AT Score), **C.** Confidence Shift between final and initial estimation*. The raincloud plots represent raw data as individual data points per participant and an overall response distribution pattern across social information proximity conditions. The black rings indicate the mean values.

**Figure 3.**
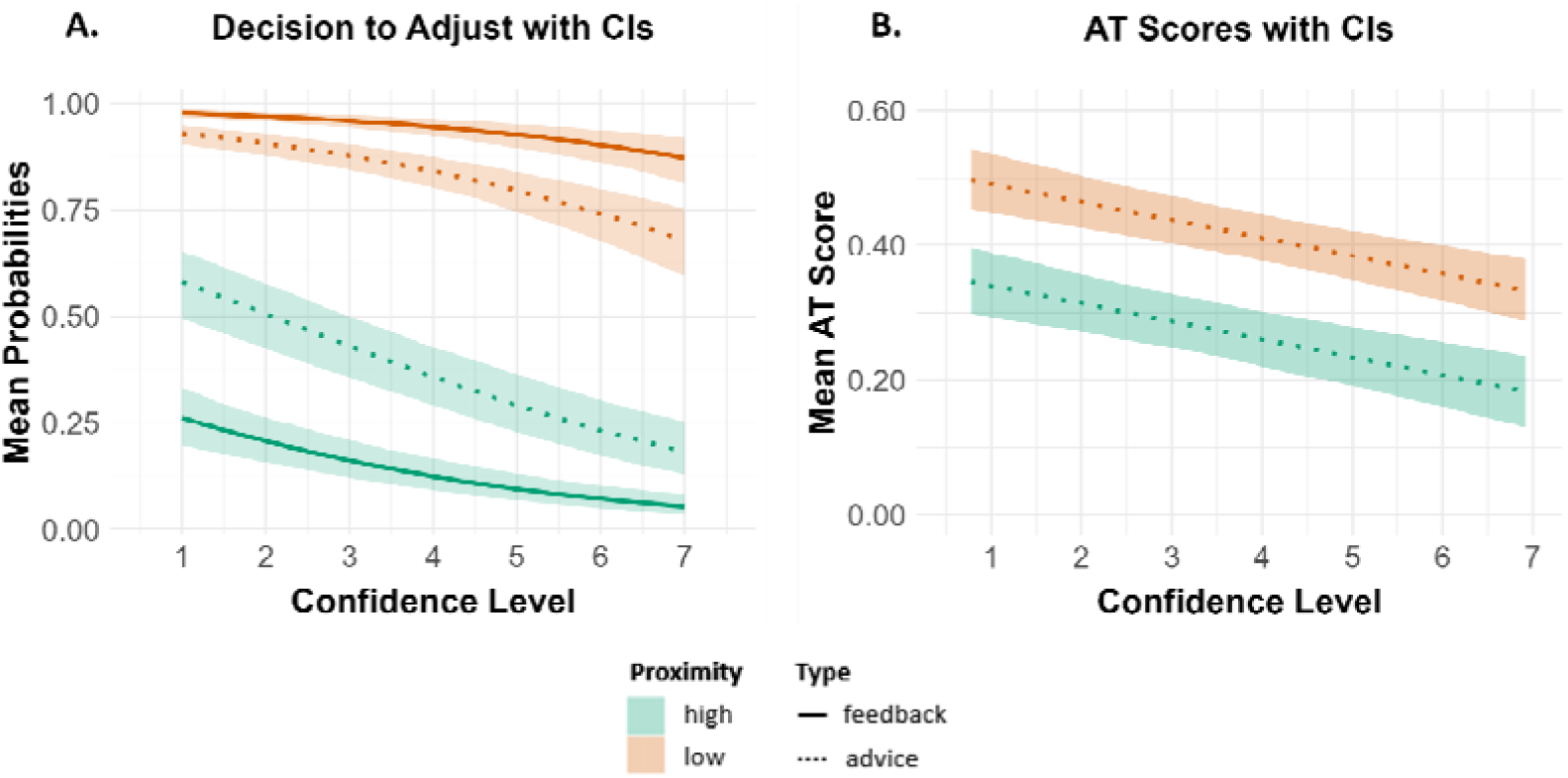
*Effect of Initial Confidence on **A.** Decision to Adjust **B**. AT Score (only advice).* The lines indicate mean bootstrapped values for the 1 to 7 confidence levels, along with their shaded 95 % confidence intervals (CIs).

***Relative Decision Change (AT Score;*** *Figure 2B).* The AT Score was significantly affected by advice proximity (β = 0.147, SE = 0.017, t = 8.491, p < .001). Compared with high-proximity advice, low-proximity elicited greater weighting of external input. The model also revealed a significant main effect of initial confidence, indicating that reliance on advice decreased with increasing confidence (β = –0.024, SE = 0.004, t = –5.716, p < .001; *Figure 3B*).

***Confidence Shift*** *(Figure 2C).* ‘Changes in confidence (final − initial) were modulated by an interaction between proximity and information type ( β = −0.983, SE = 0.102, t = –9.666, p < .001). For advice, confidence increased more after high-than low-proximity information (β = 0.763, SE = 0.0510, z = 14.947, p < .001). For feedback, this effect was even more pronounced (β = 1.746, SE = 0.101, z = 17.128, p < .001). Importantly, low-proximity advice still produced a small confidence gain (M = 0.469, CI = [0.334, 0.605]), while low proximity feedback decreased confidence (M = −0.246, CI = [−0.337, −0.155]).

### Electrophysiological data

#### ERP amplitudes

After removing trials that activated the failsafe mechanism and those rejected during the EEG preprocessing (2,999 trials, 17 % removed), a total of 13,861 trials were included in the statistical analyses (advice, high proximity = 3,528; advice, low proximity =3,388; feedback, high proximity = 3,464; feedback, low proximity = 3,481).

***FRN.*** Single-trial FRN amplitudes (*Figure 4A1*) were modulated by an interaction between social information type and initial confidence (β = 0.375, SE = 0.108, t = 3.471, p = .001). Specifically, compared to advice (β = –0.113, CI = [–0.260, 0.0344]), FRN amplitudes for feedback became less negative (i.e., more positive) with increasing initial confidence (β = 0.262, CI = [0.107, 0.417]).

**Figure 4.**
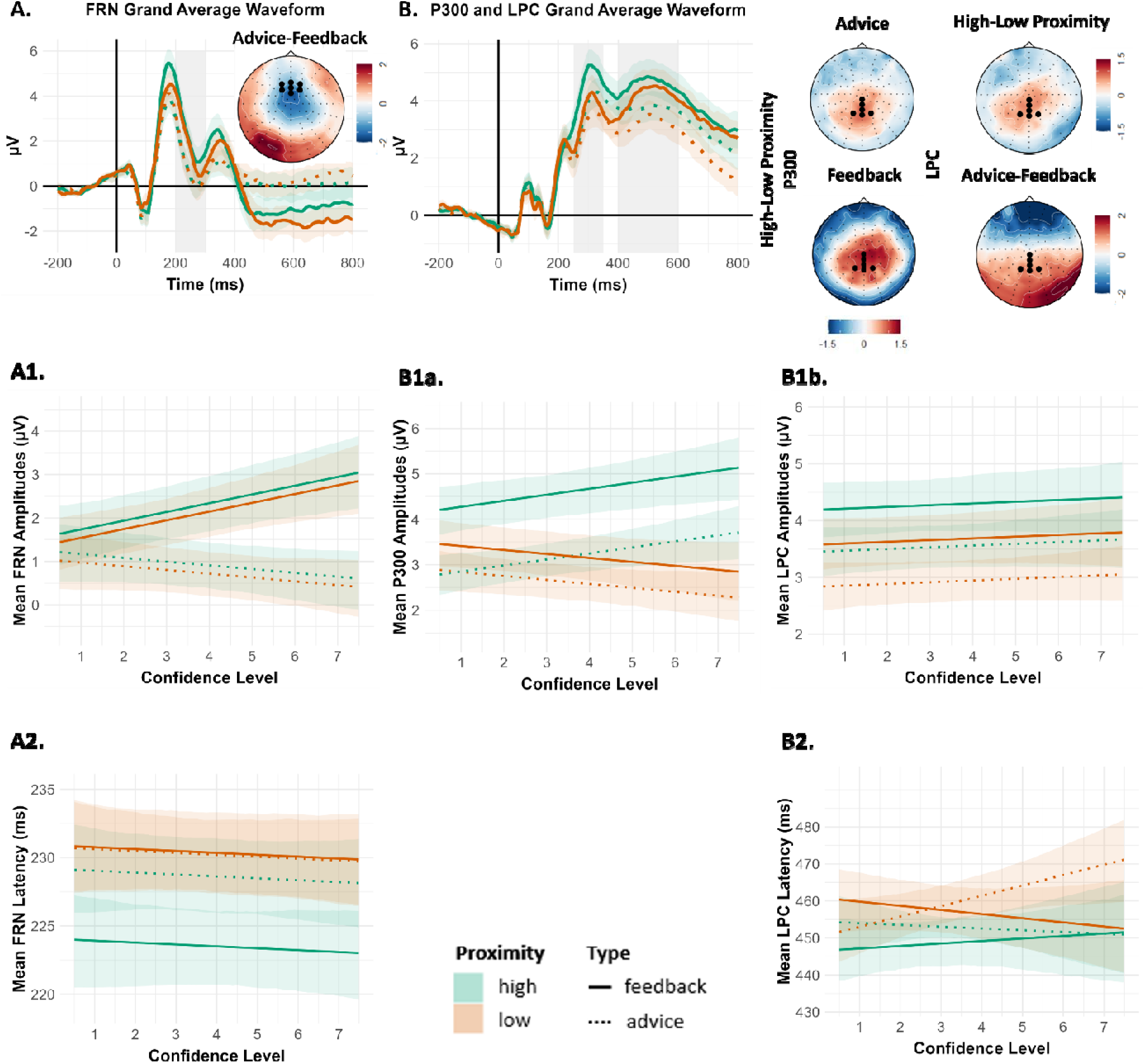
Grand average ERP waveforms of the respective ROI channels, time-locked to the presentation of social information (advice/feedback). The time window for each of the ERP components of interest is shaded in gray (***A.*** *FRN (negative going ERP), **B.** P300 and LPC)*. Topographical plots for *FRN* indicate difference maps between social information types (advice minus feedback). Topographical plots for P300 indicate difference maps between proximities (high minus low) and LPC topographical maps indicate difference maps between proximities (high minus low) at low and high initial confidence. Electrodes of interest are highlight d as black dots on the head maps. Mean bootstrapped ERP amplitudes are plotted as a function of initial confidence (***A1.*** *FRN, **B1a.** P300, **B1b.** LPC )* along with their shaded 95 % confidence intervals (CIs). Mean bootstrapped latency of ERP onsets are plotted as a function of initial confidence (***A2.*** *FRN, **B2.** LPC)* along with their shaded 95 % CIs.

***P300.*** Single-trial P300 amplitudes *(Figure 4B1a)* were modulated by an interaction between social information proximity and initial confidence (β = 0.287, SE = 0.083, t = 3.461, p = .001). P300 amplitudes increased with increasing initial confidence for high-proximity information (β = 0.173, CI = [0.064, 0.359]) but not for low-proximity social information (β = –0.114, CI = [– 0.236, 0.009])). In addition, there was a significant interaction of social information proximity and social information type (β = 0.853, SE = 0.183, t = 4.656, p <.001). That is, compared to the proximity effects for advice (β = 0.475, SE =0.112, z = 4.232, p < .001), they are more pronounced for feedback (β = 1.328, SE = 0.161, z = 8.251, p <.001).

***LPC.*** Single-trial LPC amplitudes *(Figure 4B1b)* showed main effects of both social information proximity and social information type (proximity: β = –0.615, SE = 0.096, t = –6.380, p < .001; type: β = 0.741, SE = 0.176, t = 4.213, p < .001). LPC amplitudes were reduced for low-compared to high-proximity social information and were overall higher for feedback than for advice. There was no effect of initial confidence (β = 0.040, SE = 0.054, t = 0.730, p = .469).

#### ERP Onset Latencies

***FRN.*** FRN onset latencies *(Figure 4A2)* were modulated by an interaction of social information proximity and social information type (β = 5.265, SE = 1.500, t = 3.510, p = .004). FRN onsets were delayed for low-compared to high-proximity feedback (β = 6.845, SE = 1.178, z = 5.813, p < .001), whereas no significant proximity difference was observed for advice (β = 1.580 SE = 0.918, z = 1.721, p = .312).

***P300.*** The model for P300 onset latencies did not reveal a significant full-null model comparison and was therefore not evaluated further.

***LPC.*** The LPC onset latencies *(Figure 4B2.)* were modulated by a three-way interaction between social information proximity, type, and initial confidence (β = –6.567, SE = 3.029, t = –2.168, p = .034). With increasing initial confidence, LPC onsets for advice tended to increase for low-compared to high-proximity (β = 4.241, SE = 2.10, z = 2.016, p = .182), whereas for feedback, the opposite pattern emerged, with visually slightly earlier onsets for low-compared to high-proximity feedback (β = –2.325, SE = 2.22, z = –1.047, p = .721). For the contrasts within proximity, low-proximity advice induced slower onsets with increasing confidence compared to low-proximity feedback (β = 5.080, SE = 2.17, z = 2.336, p = .09), whereas there was no difference for high-proximity advice compared to high-proximity feedback (β = −1.487, SE = 2.18, z = −0.682, p = .90).

## Discussion

Social information can guide behavior through two separable routes: an evaluative route that determines *whether* to revise one’s judgment and an informational route that specifies *how* to revise it. By combining behavioral and electrophysiological measures, the present study demonstrates that these routes unfold in distinct temporal windows for feedback and advice. Feedback engaged early evaluative mechanisms reflected in the FRN, whereas advice induced delayed confidence-dependent LPC onset latencies indicating deliberate integration of external input. Together, the data reveal that the brain distinguishes *being judged* from *being informed* – a dissociation that depends on both the proximity of social information and an individual’s confidence in their initial estimate.

The behavioral results replicated and extended previous work from the advice-taking literature (Yaniv, 2004a; Schultze et al., 2015; Du et al., 2019). Low-proximity social information increased both the likelihood of adjustment (*Decision to Adjust*) and the weighting of external input (*AT Score*; Schultze et al., 2015; Du et al., 2019). For feedback, proximity effects were stronger than for advice, reflected in high adjustment rates following low-proximity feedback even at high confidence (around 80%) and low adjustment rates after high-proximity feedback at low confidence (around 25%). In contrast, proximity effects for advice were more moderate: participants continued to adjust even after receiving high-proximity advice, indicating that advice provides not only evaluation but also guidance about the magnitude of change. Because advice is more ambiguous, individuals may balance their own and external estimates even when agreement is high, reducing proximity contrasts overall.

Feedback carries heightened motivational salience because it explicitly conveys success or failure, engaging self-referential evaluation alongside task processing (Kluger & DeNisi, 1996; Hattie & Timperley, 2007). Advice, by contrast, is only indirectly evaluative; participants recognize that an advisor may provide an estimate without necessarily knowing the correct answer, rendering advice less self-relevant. This difference emerged in the confidence data: high-proximity feedback increased confidence, whereas low-proximity feedback reduced it. For advice, even low-proximity information increased confidence after revision – consistent with the idea that combining external input with one’s own judgment reduces uncertainty and perceived error (Budescu, 2006; Harvey & Fischer, 1997; Yaniv et al., 2009). Thus, feedback exerts stronger motivational effects, whereas advice primarily serves as informational enrichment, supporting a sense of control and certainty during judgment revision.

This behavioral dissociation is also important for current theorizing on socially induced belief revision. Early advice taking research already speculated on a distinction between the evaluation and integration of socially provided information (Harvey, Harries, & Fischer, 2000), and this distinction is also reflected in sophisticated approaches such as Himmelstein’s (2022) *dual-hurdle model* of decision revision. Consistent with these approaches, our results provide evidence that individuals first decide *whether* to revise and then *how much* to revise. Our ERP data extend this framework by showing that these cognitive stages are temporally dissociable at the neural level. The FRN reflects the early evaluative hurdle, varying with confidence and proximity within feedback, whereas later components, especially LPC onset dynamics, seem to correspond to the informational hurdle engaged during advice-based integration.

Consistent with prior work showing that social influence decreases as confidence increases (Pescetelli & Yeung, 2021; Desender et al., 2018; Frömer et al., 2021), a main effect of confidence was evident behaviorally. This pattern was paralleled in the neural data showing a modulation of the FRN by an interaction between information type and confidence. Early performance monitoring revealed diminishing FRN amplitudes with increasing confidence for feedback, consistent with reinforcement-learning and cognitive-control accounts in which confidence tunes performance-monitoring signals (Braem & Egner, 2018; Holroyd et al., 2009). Advice, in contrast, did not show this confidence dependence, implying that advice does not trigger the same reflexive evaluation of outcome valence. Rather, advice engages sustained comparison and interpretation processes even at this early stage. These results suggest that confidence calibrates evaluative monitoring primarily for feedback, whereas advice processing involves more sustained comparison independent of confidence. FRN onset latencies further support this dissociation, showing delayed responses for low-versus high-proximity feedback but no latency differences for advice. The absence of proximity effects on the FRN contrasts with prior findings by Du et al. (2019), likely due to our limited deviation range and absence of a 0%-baseline. Moreover, the lack of asymmetrical outcome probabilities or explicit incentives – factors known to enhance FRN amplitude (Holroyd et al., 2009) – may have attenuated sensitivity to proximity-related expectancy violations. Our findings dovetail with recent work demonstrating separable neural signatures of reward and affective prediction errors (Heffner et al., 2025), further supporting a temporal architecture that distinguishes rapid evaluative feedback processing from later integrative stages of social information use.

Both proximity and confidence shape social information processing most prominently at later stages (Frömer et al., 2021; Zheng et al., 2021). This is reflected in the P300, which indexed the integration of confidence-based expectations with the motivational salience of social information proximity. Consistent with the social-validation effect (Schultze et al., 2015; Wanzel et al., 2017), P300 amplitudes were enhanced when social information was of high proximity, confirming individuals’ judgments. Confidence likely acted as a proxy for outcome expectation, sharpening the perceived match or mismatch between expectation and the outcome. This pattern aligns with findings that P300 amplitudes increase when outcomes match expectations and decrease when they violate them (Wu & Zhou, 2009; Rollwage et al., 2020), and with the idea that social endorsement itself conveys affective reassurance (Gibbons et al., 2003). Notably, proximity effects on P300 were more pronounced for feedback than advice, reflecting the higher motivational relevance and categorical nature of evaluative feedback (Kluger & DeNisi, 1996; Hattie & Timperley, 2007; Belschak & Den Hartog, 2009), whereas advice primarily engages P300 through its informational relevance.

Contrary to our initial hypotheses, LPC amplitudes primarily reflected the social validation effects of the high proximity social information, paralleling the P300 amplitude effects rather than signaling increased processing of low-proximity information. This suggests that the LPC integrates the extracted motivational relevance of social information with evaluative updating (Herbert et al., 2006; Schacht et al., 2012; Grassi et al., 2023; Ziereis & Schacht, 2024), resulting in overall larger amplitudes for feedback than advice. While LPC amplitudes themselves did not vary with confidence, onset latencies showed distinct patterns for feedback and advice. For feedback, low proximity delayed LPC onsets when confidence was low, but this delay was reduced as confidence increased – consistent with reduced deliberation when participants felt certain, paralleling the reduced FRN amplitudes at higher confidence. In contrast, advice elicited progressively slower LPC onsets for low-proximity trials as confidence increased, indicating intensified deliberation when confident individuals encountered discrepant advice. Thus, for advice, confidence sharpens expectation–outcome mismatches that guide the reconciliation between external input and one’s own judgment.

Although our findings delineate distinct temporal routes of social information processing, several limitations warrant consideration. The interpretation of proximity for advice may vary across individuals: for some, a difference of 10% may appear reassuringly close, for others already discrepant. Moreover, we focused on confidence as a key moderator, but previous research emphasizes that perceived advisor expertise also critically determines informational influence (Harvey & Fischer, 1997; Yaniv & Kleinberger, 2000; Meshi et al., 2012). Future work should therefore examine how confidence and perceived expertise jointly shape neural and behavioral sensitivity to social information. Finally, while the present design allowed precise temporal inference under controlled conditions, testing whether the same dissociations hold in more naturalistic, dynamic interactions remains an important step (Falk & Bassett, 2017).

At a functional level, our data clarify why feedback and advice, though closely related, serve complementary roles in judgment and decision making. Feedback provides categorical evaluative information – deciding *whether* an initial judgment was correct – whereas advice adds directional information that guides *how much* to adjust, albeit at the cost of a more equivocal evaluative component. The FRN reflects confidence-weighted evaluative monitoring tuned to outcome relevance, while the P300 and LPC capture the motivational and deliberative integration of social input with self-generated certainty. Confidence sharpened the reward value of confirmatory feedback but also intensified deliberation when confident individuals confronted discrepant advice. Behaviorally, while their confidence influenced participants’ decision to adjust their estimates, these mechanisms yielded pronounced proximity effects for feedback but more moderate effects for advice. At a mechanistic level, feedback engages confidence-weighted performance monitoring (FRN), while advice triggers sustained deliberative integration (LPC dynamics). The P300 bridges these processes, integrating the motivational salience of social validation with task-relevant updating.

Together, these findings advance a mechanistic account of how metacognitive and social processes jointly determine judgment revision. They reveal that humans arbitrate between internal certainty and external input through two temporally coordinated routes – an evaluative route determining *whether* to revise and an informational route specifying *how* to revise. This dual-route architecture has implications for education (pairing evaluative feedback with concrete guidance), collective decision-making (harnessing diverse perspectives), human-AI collaboration (providing both confidence signals and actionable recommendations), and misinformation research (explaining why confidence shields against superficial pressure yet preserves consideration of substantive arguments). By situating confidence and proximity as key computational variables, this work offers a framework for optimizing social influence across contexts from education to human-AI cooperation.

## Acknowledgements

This project was funded by an Audacity Fund from the Leibniz ScienceCampus Primate Cognition (LSC-AF2020_04) and by the Deutsche Forschungsgemeinschaft (DFG, Project-ID 454648639 - SFB 1528). The authors would like to thank all the student assistants of the lab for helping with data collection. They would also like to acknowledge Dr. Francesco Grassi for programming the experiment in Python, Patrick Sautner and Esther Semmelhack for their assistance in pre-processing the EEG data in MATLAB and lastly Dr. Roger Mundry for providing functions for statistical analyses.

## Data availability

The code of this study is available upon request from the corresponding author SM. The pre-processed EEG data is available on the Open Science Framework (https://osf.io/nvfh5). Raw EEG data is available upon request to the authors. Parts of the data of this study were presented at conferences and scientific meetings.

## CRediT Authorship Contribution Statement

Sriranjani Manivasagam: Investigation, Data curation, Formal analysis, Visualization, Writing – original draft, Writing – review & editing. Thomas Schultze: Conceptualization, Methodology, Validation, Writing – review & editing. Anne Schacht: Conceptualization, Methodology, Validation, Supervision, Project administration, Funding acquisition, Writing – review & editing. Anna Fischer: Methodology, Investigation, Supervision, Validation, Writing – review & editing

