## Supplementary Materials for "Distinct neural routes for evaluative and informational social influence"

Both the behavioral and EEG models have 'advice' and 'high' as the reference levels for the factors 'social information type' (Soc.Info.Type) and 'social information proximity' (Soc.Info.Prox) respectively. An exception was made for the models predicting P300 amplitudes, where the level 'low' was used as the reference for 'social information type'. For the behavioral models, the trials are coded within each 'advice' and 'feedback' block (1-100) and was z transformed (mean:50.856; sd:28.724). For the EEG models, the trials are coded within each combination of 'social information type' and 'social information proximity' (1-50) and was z transformed (mean:24.905; sd:14.053). Confidence (Initial confidence) ratings (1–7) were z transformed within subject to control for individual differences in scale use. Thus, for each participant, a value of 0 reflects their average confidence, positive values reflect above-average confidence, and negative values reflect below-average confidence. Because scaling was done per subject, the transformed values cannot be directly back-transformed to a global mean and sd. Partial $R^2$ was calculated using the method suggested by Nakagawa & Schielzeth (2013) along with their confidence intervals was calculated using the 'r2beta' function of the package r2glmm (version 0.1.2; Jaeger et al., 2017).

**Notes:** β = model estimate, SE = standard error of the estimate, CI = lower and upper 95% bootstrapped confidence intervals, Stab = estimate ranges leaving out one participant at a time, pSatter = Satterwhite approximated p-values, $R^2$ partial = partial $R^2$ with lower and upper confidence intervals, LRT = Likelihood ratio test.

p-values of the intercepts not shown due to limited interpretability.

#### Behavioral models results table

##### 1. Decision to Adjust

**Formula:** Dec_to_adjust ~ Soc.Info.Type*Soc.Info.Prox + Conf1 + Trial + (1 + Soc.Info.Type*Soc.Info.Prox*Conf1 + Trial|sub_id)

| | estimate | std. error | t value | p value | stab min | stab max | lower CI | upper CI |
|---|---|---|---|---|---|---|---|---|
| (Intercept) | -0.251 | 0.166 | -1.510 | - | -0.300 | -0.220 | -0.589 | 0.084 |
| Soc. Info. Type feedback | -1.434 | 0.213 | -6.746 | <.001 | -1.494 | -1.364 | -1.828 | -1.037 |
| Soc. Info. Prox low | 2.210 | 0.127 | 17.340 | <.001 | 2.180 | 2.251 | 1.969 | 2.455 |
| Initial confidence | -0.277 | 0.036 | -7.718 | <.001 | -0.288 | -0.267 | -0.352 | -0.206 |
| Trial | -0.109 | 0.029 | -3.709 | <.001 | -0.119 | -0.101 | -0.169 | -0.053 |
| Soc. Info. Type feedback : Soc. Info. Prox low | 2.697 | 0.270 | 9.975 | <.001 | 2.627 | 2.777 | 2.159 | 3.234 |

**Table 1:** The model's overall R2 partial was: .148 [.158, .139]. Likelihood Ratio tests indicated significant contribution of the interaction between Soc. Info.Type and Soc. Info.Prox (Chisq = 69.751, p <.001) and Conf1 (Chisq = 45.620, p < .001).

##### 2. AT Score

**Formula:** AT Score ~ Soc.Info.Prox + Conf1 + Trial + (1 + Soc.Info.Prox*Conf1 + Trial|sub_id)

| | estimate | std. error | t value | p value | stab min | stab max | lower CI | upper CI |
|---|---|---|---|---|---|---|---|---|
| (Intercept) | 0.291 | 0.022 | 12.976 | - | 0.284 | 0.295 | 0.246 | 0.337 |
| Soc. Info. Prox low | 0.147 | 0.017 | 8.491 | <.001 | 0.143 | 0.153 | 0.114 | 0.182 |
| Initial confidence | -0.024 | 0.004 | -5.716 | <.001 | -0.025 | -0.023 | -0.033 | -0.016 |
| Trial | -0.015 | 0.005 | -2.897 | .005 | -0.017 | -0.013 | -0.025 | -0.005 |

**Table 2:** The model's overall R2 partial was: .045 [.055,.037]. Likelihood Ratio tests indicated significant contribution of Soc. Info.Prox (Chisq = 52.266, p <.001) and Conf1 (Chisq = 27.588, p < .001).

##### 3. Confidence Shift

Formula: Conf_Shift ~ Soc.Info.Type*Soc.Info.Prox + Trial + (1 + Soc.Info.Type*Soc.Info.Prox + Trial|sub_id)

| | estimate | std. error | t value | p value | stab min | stab max | lower CI | upper CI |
|---|---|---|---|---|---|---|---|---|
| (Intercept) | 1.232 | 0.072 | 17.140 | - | 1.206 | 1.247 | 1.093 | 1.367 |
| Soc. Info. Type feedback | 0.267 | 0.073 | 3.653 | <.001 | 0.246 | 0.290 | 0.125 | 0.412 |
| Soc. Info. Prox low | -0.763 | 0.051 | -14.947 | <.001 | -0.775 | -0.744 | -0.863 | -0.657 |
| Trial | 0.035 | 0.015 | 2.353 | .022 | 0.030 | 0.040 | 0.005 | 0.065 |
| Soc. Info. Type feedback : Soc. Info. Prox low | -0.983 | 0.102 | -9.666 | <.001 | -1.007 | -0.944 | -1.178 | -0.773 |

**Table 3:** The model's overall R2 partial was: .240 [.251,.230]. Likelihood Ratio tests indicated significant contribution of the interaction between Soc. Info.Type and Soc. Info.Prox (Chisq = 63.426, p <.001).

#### Electrophysiological data results table

##### 4. Feedback Related Negativity (FRN)

Formula: FRN ~ Soc.Info.Prox + Soc.Info.Type*Conf1 + Trial + (1+ Soc.Info.Type*Soc.Info.Prox *Conf1 + Trial|sub_id)

| | estimate | std. error | t value | p value | stab min | stab max | lower CI | upper CI |
|---|---|---|---|---|---|---|---|---|
| (Intercept) | 0.984 | 0.267 | 3.692 | - | 0.898 | 1.080 | 0.455 | 1.530 |
| Soc. Info. Prox low | -0.193 | 0.103 | -1.877 | .063 | -0.238 | -0.168 | -0.395 | 0.003 |
| Soc. Info. Type feedback | 1.177 | 0.217 | 5.423 | <.001 | 1.089 | 1.233 | 0.727 | 1.587 |
| Initial confidence | -0.113 | 0.075 | -1.502 | .141 | -0.131 | -0.093 | -0.256 | 0.035 |
| Trial | -0.087 | 0.056 | -1.555 | .125 | -0.100 | -0.068 | -0.198 | 0.018 |
| Soc. Info. Type feedback : Initial confidence | 0.375 | 0.108 | 3.471 | <.001 | 0.341 | 0.402 | 0.152 | 0.586 |

**Table 4:** The model's overall R2 partial was: .011 [.015,.008]. Likelihood Ratio tests indicated significant contribution of the interaction between Soc. Info.Type and Initial confidence (Chisq = 10.960, p =.001).

## 5. P300

Formula: P300 ~ Soc.Info.Type*Soc.Info.Prox + Soc.Info.Prox*Conf1 + Trial + (1 + Soc.Info.Type*Soc.Info.Prox *Conf1 + Trial|sub_id)

| | estimate | std. error | t value | p value | stab min | stab max | lower CI | upper CI |
|---|---|---|---|---|---|---|---|---|
| (Intercept) | 2.656 | 0.194 | 13.683 | - | 2.595 | 2.720 | 2.255 | 3.023 |
| Soc. Info. Type feedback | 0.571 | 0.198 | 2.890 | .006 | 0.510 | 0.606 | 0.168 | 0.979 |
| Soc. Info. Prox high | 0.475 | 0.112 | 4.232 | <.001 | 0.448 | 0.507 | 0.239 | 0.717 |
| Initial confidence | -0.114 | 0.063 | -1.814 | .095 | -0.138 | -0.090 | -0.235 | 0.010 |
| Trial | -0.003 | 0.042 | -0.065 | .966 | -0.014 | 0.010 | -0.093 | 0.080 |
| Soc. Info. Type feedback : Soc. Info. Prox high | 0.853 | 0.183 | 4.656 | <.001 | 0.813 | 0.900 | 0.483 | 1.215 |
| Soc. Info. Prox high : Initial confidence | 0.287 | 0.083 | 3.461 | .001 | 0.265 | 0.323 | 0.130 | 0.447 |

**Table 5:** The model's overall R2 partial was: .021 [.026,.017]. Likelihood Ratio tests indicated significant contribution of the interaction between Soc. Info.Type and Initial confidence (Chisq = 9.331, p =.002) and the interaction between Soc. Info.Type and Soc. Info.Prox (Chisq = 19.403, p <.001).

#### 6. LPC

Formula: LPC ~ Soc.Info.Type + Soc.Info.Prox + Conf1 + Trial + (1 + Soc.Info.Type*Soc.Info.Prox*Conf1* Trial|sub_id)

| | estimate | std. error | t value | p value | stab min | stab max | lower CI | upper CI |
|---|---|---|---|---|---|---|---|---|
| (Intercept) | 3.535 | 0.194 | 18.221 | - | 3.461 | 3.610 | 3.172 | 3.905 |
| Soc. Info. Type feedback | 0.741 | 0.176 | 4.213 | <.001 | 0.603 | 0.774 | 0.404 | 1.112 |
| Soc. Info. Prox low | -0.615 | 0.096 | -6.380 | <.001 | -0.657 | -0.581 | -0.793 | -0.426 |
| Initial confidence | 0.040 | 0.054 | 0.730 | .469 | 0.023 | 0.053 | -0.062 | 0.147 |
| Trial | 0.056 | 0.050 | 1.105 | .271 | 0.043 | 0.066 | -0.046 | 0.156 |

**Table 6:** The model's overall R2 partial was: .008 [.012,.006]. Likelihood Ratio tests indicated significant contribution of the Soc. Info.Type (Chisq = 12.599, p <.001) Soc. Info.Prox (Chisq = 30.414, p <.001).

#### 7. FRN onset latency

Formula: Onset Latency ~ Soc.Info.Type*Soc.Info.Prox + Conf1 + Trial + (1+ Soc.Info.Type*Soc.Info.Prox*Conf1 + Trial|sub_id)

| | estimate | std. error | t value | p value | stab min | stab max | lower CI | upper CI |
|---|---|---|---|---|---|---|---|---|
| (Intercept) | 228.747 | 1.355 | 168.766 | - | 228.150 | 229.049 | 226.141 | 231.517 |
| Soc. Info. Type feedback | -5.121 | 1.311 | -3.905 | .002 | -5.370 | -4.403 | -7.821 | -2.786 |
| Soc. Info. Prox low | 1.580 | 0.918 | 1.721 | .244 | 1.388 | 1.963 | -0.131 | 3.255 |
| Initial confidence | -0.180 | 0.356 | -0.507 | .943 | -0.225 | 0.105 | -0.841 | 0.492 |
| Trial | 0.427 | 0.359 | 1.188 | .195 | 0.347 | 0.541 | -0.311 | 1.150 |
| Soc. Info. Type feedback : Soc. Info. Prox low | 5.265 | 1.500 | 3.510 | .004 | 4.587 | 5.700 | 2.317 | 8.452 |

**Table 7:** The model's overall R2 partial was: .005 [.009,.003]. Likelihood Ratio tests indicated significant contribution of the interaction between Soc. Info.Type and Soc. Info.Prox (Chisq = 78.364, p <.001).

#### 8. P300 onset latency

| | npar | AIC | BIC | loglik | deviance | chisq | df | p value |
|---|---|---|---|---|---|---|---|---|
| Null model | 48 | 98689.73 | 99035.27 | -49296.87 | 98593.73 | - | - | - |
| Full model | 51 | 98888.68 | 99255.81 | -49393.34 | 98786.68 | 0 | 3 | 1 |

**Table 8:** Full-Null comparison table

#### 9. LPC onset latency

Formula: Onset Latency ~ Soc.Info.Type*Soc.Info.Prox*Conf1 + Trial + (1 + Soc.Info.Type*Soc.Info.Prox*Conf1 + Trial|sub_id)

| | estimate | std. error | t value | p value | stab min | stab max | lower CI | upper CI |
|---|---|---|---|---|---|---|---|---|
| (Intercept) | 452.976 | 1.380 | 328.319 | - | 452.533 | 454.129 | 449.999 | 455.531 |
| Soc. Info. Type feedback | -4.402 | 3.401 | -1.294 | .201 | -5.319 | -3.795 | -10.850 | 2.648 |
| Soc. Info. Prox low | 5.950 | 2.989 | 1.991 | .050 | 5.376 | 6.625 | 0.639 | 12.193 |
| Initial confidence | -0.614 | 1.648 | -0.373 | .708 | -1.137 | -0.089 | -3.738 | 2.698 |
| Trial | 0.271 | 0.736 | 0.368 | .715 | 0.071 | 0.463 | -1.195 | 1.726 |
| Soc. Info. Type feedback : Soc. Info. Prox low | 2.906 | 3.918 | 0.742 | .462 | 2.046 | 3.701 | -5.240 | 10.179 |
| Soc.Info.Typefeedback:Confidence1 | 1.487 | 2.181 | 0.682 | .495 | 0.698 | 2.142 | -2.908 | 5.776 |
| Soc.Info.Proxlow:Confidence1 | 4.241 | 2.104 | 2.016 | .046 | 3.389 | 4.715 | 0.155 | 8.262 |
| Soc. Info. Type feedback : Soc. Info. Prox low : Initial confidence | -6.567 | 3.029 | -2.168 | .034 | -7.300 | -5.210 | -12.165 | -0.565 |

**Table 9:** The model's overall R2 partial was: .004 [.007,.002]. Likelihood Ratio tests indicated significant contribution of the interaction between Soc. Info.Type and Soc. Info.Prox and Initial Confidence (Chisq = 4.484, p = .034).

#### Pre-registered models

For the pre-registered models, both the behavioral and EEG models have 'advice' and 'high' as the reference levels for the factors 'social information type' (Soc.Info.Type) and 'social information proximity' (Soc.Info.Prox) respectively.

#### Results

#### Behavioral

***Decision to Adjust.*** 'Decision to Adjust' was quantified as a binary variable (0/1), which captured the probability with which participants opted to revise their estimates after receiving social information. It was modulated by an interaction between social information proximity and social information type (proximity x type: $\beta = 2.634$, SE = 0.257, z = 10.256, $p < .001$). While participants adjusted their initial estimates more often after receiving advice of low compared to high proximity (contrast $_{(advice\_low-advice\_high)}$: $\beta = 2.07$, SE = 0.123, z = 16.831, $p < .001$), for feedback, this difference was further pronounced (contrast $_{(feedback\_low-feedback\_high)}$: $\beta = 4.71$, SE = 0.220, z = 21.410, $p < .001$).

***AT Score.*** To calculate 'AT Score', values bigger than one and smaller than zero were winsorized to one and zero respectively. The overall percentage of trials that were winsorized was 10.43 %. 'AT Score' was modulated by advice proximity (proximity: $\beta = 0.145$, SE = 0.017, t = 8.386, $p < .001$). Compared to the high proximity advice, low proximity advice led to a significantly greater 'AT Score', that is, advice was relatively weighted more when it was low in proximity to the participant's initial estimate.

***Confidence Shift.*** 'Confidence Shift' was modulated by an interaction between social information proximity and social information type (proximity x type: $\beta = -0.983$, SE = 0.102, t = - 9.632, $p < .001$). For advice, high proximity led to larger confidence gains compared to low proximity (contrast $_{(advice\_high - advice\_low)}$: $\beta = 0.760$, SE = 0.0513, z = 14.818, $p < .001$). For feedback, the effect was more pronounced, with high proximity resulting in a substantially greater increase in confidence compared to low proximity (contrast $_{(feedback\_high - feedback\_low)}$: $\beta = 1.743$, SE = 0.101, z = 17.135, $p < .001$). Notably, while low proximity advice still increased confidence (M = 0.469, CI = [0.334, 0.605]), low proximity feedback decreased confidence (M = -0.248, CI = [-0.339, -0.156]).

**ERP Amplitudes**

***FRN.*** FRN single-trial amplitudes were not modulated by social information proximity (proximity: $\beta = -0.154$, SE = 0.097, t = -1.587, p = .116). On the contrary, the model indicated a significant main effect of social information type, where feedback had reduced negative (more positive) amplitudes compared to advice (type: $\beta = 1.211$, SE = 0.214, t = 5.661, $p < .001$).

***P300.*** P300 single-trial amplitudes were modulated by an interaction between social information proximity and social information type (proximity x type: $\beta = 0.675$, SE = 0.177, t = 3.803, $p < .001$), Specifically, for advice, the P300 amplitudes were enhanced for high proximity compared to the low proximity advice (contrast $_{(advice\_high - advice\_low)}$: $\beta = 0.466$, SE = 0.104, t = 4.462, $p <.001$), while for feedback we found a more pronounced enhancement of the P300 amplitudes for high compared to low proximity feedback (contrast $_{(feedback\_high - feedback\_low)}$: $\beta = 1.141$, SE = 0.164, t = 6.956, $p <.001$).

***LPC.*** LPC single-trial amplitudes were modulated by a main effect of social information proximity, where low proximity social information had reduced LPC amplitude compared to high (proximity: β = -0.576, SE = 0.099, t = -5.823, p <.001). Additionally, the model indicated a significant main effect of social information type, where feedback had enhanced LPC amplitudes compared to advice (type: β = 0.766, SE = 0.185, t = 4.148, p < .001).

#### ERP Onset Latencies

The FRN onset latencies were modulated by social information proximity, where low proximity conditions had significantly slower mean onsets in comparison to the high proximity conditions (proximity: β = 3.541, SE = 0.694, t = 5.099, p < .001). The P300 onset latencies were modulated by social information type, showing overall faster onsets for feedback compared to advice (type: β = - 3.703, SE = 1.058, t = - 3.499, p = .001. The LPC onset latencies were modulated by a main effect of proximity, where low proximity had an overall slower onset compared to high (proximity: β = 7.186, SE = 1.964, t = 3.658, p <.001).

**Notes:** β = model estimate, SE = standard error of the estimate, CI = lower and upper 95% bootstrapped confidence intervals, Stab = estimate ranges leaving out one participant at a time, pSatter = Satterwhite approximated p-values, $R^2$ partial = partial $R^2$ with lower and upper confidence intervals, LRT = Likelihood ratio test.

p-values of the intercepts not shown due to limited interpretability.

#### Pre-registered behavioral models results table

##### 1. Decision to Adjust

| | estimate | std. error | t value | p value | stab min | stab max | lower CI | upper CI |
|---|---|---|---|---|---|---|---|---|
| (Intercept) | -0.200 | 0.166 | -1.206 | - | -0.243 | -0.168 | -0.534 | 0.107 |
| Soc. Info. Type feedback | -1.404 | 0.209 | -6.733 | <.001 | -1.456 | -1.344 | -1.789 | -0.989 |
| Soc. Info. Prox low | 2.072 | 0.123 | 16.831 | <.001 | 2.042 | 2.105 | 1.833 | 2.309 |
| Soc. Info. Type feedback : Soc. Info. Prox low | 2.634 | 0.257 | 10.256 | <.001 | 2.571 | 2.692 | 2.145 | 3.128 |

**Table 10:** The model's overall R2 partial was: .153 [.163,.143]. Likelihood Ratio tests indicated significant contribution of the int eraction between Soc. Info.Type and Soc. Info.Prox (Chisq = 72.525, p <.001).

#### 2. AT Score

| | estimate | std. error | t value | p value | stab min | stab max | lower CI | upper CI |
|---|---|---|---|---|---|---|---|---|
| (Intercept) | 0.291 | 0.022 | 13.120 | - | 0.284 | 0.294 | 0.245 | 0.332 |
| Soc. Info. Prox low | 0.145 | 0.017 | 8.386 | <.001 | 0.141 | 0.150 | 0.112 | 0.177 |

**Table 11:** The model's overall R2 partial was: .039 [.047,.031]. Likelihood Ratio tests indicated significant contribution of the Soc. Info.Prox (Chisq = 51.592, p <.001).

#### 3. Confidence Shift

| | estimate | std. error | t value | p value | stab min | stab max | lower CI | upper CI |
|---|---|---|---|---|---|---|---|---|
| (Intercept) | 1.229 | 0.071 | 17.294 | - | 1.205 | 1.244 | 1.092 | 1.372 |
| Soc. Info. Type feedback | 0.266 | 0.072 | 3.677 | <.001 | 0.245 | 0.289 | 0.125 | 0.402 |
| Soc. Info. Prox low | -0.760 | 0.051 | -14.818 | <.001 | -0.772 | -0.741 | -0.869 | -0.662 |
| Soc. Info. Type feedback : Soc. Info. Prox low | -0.983 | 0.102 | -9.632 | <.001 | -1.007 | -0.944 | -1.185 | -0.779 |

**Table 12:** The model's overall R2 partial was: .240 [.251,.229]. Likelihood Ratio tests indicated significant contribution of the interaction between Soc. Info.Type and Soc. Info.Prox (Chisq = 63.103, p <.001).

### Electrophysiological data

#### 4. FRN

| | estimate | std. error | t value | p value | stab min | stab max | lower CI | upper CI |
|---|---|---|---|---|---|---|---|---|
| (Intercept) | 0.905 | 0.265 | 3.418 | - | 0.806 | 1.001 | 0.430 | 1.438 |
| Soc. Info. Type feedback | 1.211 | 0.214 | 5.661 | <.001 | 1.121 | 1.262 | 0.743 | 1.603 |
| Soc. Info. Prox low | -0.154 | 0.097 | -1.587 | .116 | -0.187 | -0.126 | -0.344 | 0.036 |

**Table 13:** The model's overall R2 partial was: .011 [.014,.008]. Likelihood Ratio tests indicated significant contribution of the Soc. Info.Type (Chisq = 26.557, p <.001).

## 5. P300

| | estimate | std. error | t value | p value | stab min | stab max | lower CI | upper CI |
|---|---|---|---|---|---|---|---|---|
| (Intercept) | 2.658 | 0.197 | 13.469 | - | 2.592 | 2.700 | 2.284 | 3.059 |
| Soc. Info. Type feedback | 0.696 | 0.193 | 3.611 | .001 | 0.642 | 0.733 | 0.317 | 1.070 |
| Soc. Info. Prox high | 0.466 | 0.104 | 4.462 | <.001 | 0.442 | 0.500 | 0.268 | 0.671 |
| Soc. Info. Type feedback : Soc. Info. Prox high | 0.675 | 0.177 | 3.803 | <.001 | 0.636 | 0.721 | 0.329 | 1.033 |

**Table 14:** The model's overall R2 partial was: .017 [.022,.014]. Likelihood Ratio tests indicated significant contribution of the interaction between Soc. Info.Type and Soc. Info.Prox (Chisq = 13.621, p <.001).

#### 6. LPC

| | estimate | std. error | t value | p value | stab min | stab max | lower CI | upper CI |
|---|---|---|---|---|---|---|---|---|
| (Intercept) | 3.619 | 0.203 | 17.838 | - | 3.564 | 3.663 | 3.255 | 3.996 |
| Soc. Info. Type feedback | 0.766 | 0.185 | 4.148 | <.001 | 0.705 | 0.807 | 0.432 | 1.132 |
| Soc. Info. Prox low | -0.576 | 0.099 | -5.823 | <.001 | -0.600 | -0.543 | -0.777 | -0.397 |

**Table 15:** The model's overall R2 partial was: .007 [.010,.005]. Likelihood Ratio tests indicated significant contribution of the Soc. Info.Type (Chisq = 15.673, p <.001) and Soc.Info.Prox (Chisq = 29.314, p <.001).

#### 7. FRN onsets

| | estimate | std. error | t value | p value | stab min | stab max | lower CI | upper CI |
|---|---|---|---|---|---|---|---|---|
| (Intercept) | 226.806 | 1.315 | 172.440 | - | 226.536 | 227.233 | 224.229 | 229.468 |
| Soc. Info. Type feedback | -1.674 | 0.953 | -1.756 | .087 | -1.965 | -1.362 | -3.587 | 0.270 |
| Soc. Info. Prox low | 3.541 | 0.694 | 5.099 | <.001 | 3.376 | 3.753 | 2.136 | 4.953 |

**Table 16:** The model's overall R2 partial was: .003 [.005,.001]. Likelihood Ratio tests indicated significant contribution of Soc.Info.Prox (Chisq = 19.962, p <.001).

#### 8. P300 onsets

| | estimate | std. error | t value | p value | stab min | stab max | lower CI | upper CI |
|---|---|---|---|---|---|---|---|---|
| (Intercept) | 263.264 | 1.282 | 205.432 | - | 262.993 | 263.901 | 260.573 | 265.716 |
| Soc. Info. Type feedback | -3.703 | 1.058 | -3.499 | .001 | -3.972 | -3.348 | -5.916 | -1.724 |
| Soc. Info. Prox low | -0.545 | 0.750 | -0.727 | .470 | -0.691 | -0.322 | -2.031 | 1.017 |

**Table 17:** The model's overall R2 partial was: .003 [.005,.001]. Likelihood Ratio tests indicated significant contribution of the Soc. Info.Type (Chisq = 11.450, p =.001).

#### 9. LPC onsets

| | estimate | std. error | t value | p value | stab min | stab max | lower CI | upper CI |
|---|---|---|---|---|---|---|---|---|
| (Intercept) | 452.656 | 2.615 | 173.110 | - | 452.197 | 453.776 | 447.698 | 457.820 |
| Soc. Info. Type feedback | -3.438 | 2.271 | -1.514 | .136 | -4.088 | -2.699 | -7.800 | 1.286 |
| Soc. Info. Prox low | 7.186 | 1.964 | 3.658 | <.001 | 6.648 | 7.781 | 3.189 | 10.898 |

**Table 18:** The model's overall R2 partial was: .002 [.005,.001]. Likelihood Ratio tests indicated significant contribution of Soc.Info.Prox (Chisq = 12.408, p <.001).
